# Niche characteristics and potential distribution of *Thelocactus* species, a Mexican genus of globular cacti

**DOI:** 10.1101/124511

**Authors:** Alessandro Mosco

## Abstract

**Aim:** Although Mexican Cactaceae are a significative component of Mexican flora and have a relevant economic and ornamental value, the knowledge of the environmental factors characterising their niche is still quite incomplete. This study was aimed at defining the potential distribution and ecological niche of *Thelocactus* species.

**Methods:** Climatic and environmental variables constraining the distribution of *Thelocactus* species were identified by means of environmental niche models (ENM) and ordination techniques, and used to generate potential distribution maps. The constructed ENMs were compared to assess the similarities of the ecological niche of *Thelocactus* species and to know if they share the same ecological niche space.

**Results:** The distribution of *Thelocactus* species was mostly limited by a combination of two environmental factors, isothermality and precipitation of wettest quarter. The null hypothesis of the niche equivalency test was rejected for all paired comparisons between all *Thelocactus* species except between the pair *Thelocactus leucacanthus-Thelocactus hastifer*. The results of the niche similarity tests were quite varied, for some species pairs the niche similarities were higher than expected by chance, for others the null hypothesis was rejected, while in other species pairs niches were more similar than expected by chance, but only in one direction.

**Main conclusions:** The differences in habitat requirements were well documented by the significative differences in the niche ecological space as shown by the equivalency test, while the high percentage of niches that were more similar than expected by chance suggest a high degree of niche conservatism among *Thelocactus* species. The spatial predictions could serve to improve field design sampling to discover new populations, while niche characteristics could be relevant for improving preservation actions and guiding reintroduction programs for a better conservation of *Thelocactus* species.

## Introduction

Deserts cover between 26–35% of the land surface of the Earth and generally occur in a band around the world between 15–30° N and S latitude and their climate is dominated by low precipitation, generally below 250 mm yr^−1^ (Forseth, 2010). The largest desert in North America is the Chihuahuan Desert encompassing the southwestern United States, the Central Mexican Highlands, and extending south to disjunct parts of Chihuahuan vegetation in the states of Querétaro and Hidalgo. It is delimited from nearby regions by the large mountain ranges of the Sierra Madre Occidental to the west and the Sierra Madre Oriental to the east, which are responsible for its arid climate that associated to the mountainous landscape make it one of the most biologically diverse arid regions on earth with a high proportion of endemic species (Schmidt, 1986; Dinerstein *et al*., 2000). The Chihuahuan Desert is the Mexican region with the highest diversity of Cactaceae both at the National and Continental levels making it a hot spot of cactus diversity hosting the largest assemblage of cactus genera and species many of which are strictly endemic to this region (Hernández & Godinez A., 1994; Hernández & Bárcenas, 1995; Hernández *et al*., 2004). *Thelocactus*, a small genus of globular cacti, pertains to this group of taxa, being distributed from the southern disjunct part of the Chihuahuan Desert in Querétaro and Hidalgo to Texas, with only one species occurring outside the Chihuahuan Desert borders, in the Tamaulipan matorral or Tamaulipan mezquital regions. The genus *Thelocactus* is made up of about 12 species, according to the authority (Anderson, 1987; Mosco & Zanovello, 2000; Guzmán *et al*., 2007), with highly different distribution areas that are very wide as is the case for *Thelocactus bicolor*, which is found from Texas to San Luis Potosí, or very small as for *Thelocactus hastifer* or *Thelocactus lausseri*, which is known only from the type locality. Small geographic range sizes characterise rare cactus species, narrow endemics being restricted to areas smaller than 10 km^2^ representing nearly one-third of the Cactaceae from the Chihuahuan Desert (Hernández *et al*., 2010). The conservation status of *Thelocactus* species is generally good, only *T. hastifer* being listed in the IUCN Red List as endangered (Gómez-Hinostrosa, Sánchez & Guadalupe Martínez, 2013). The main threats derive from land conversion, collection, and residential development, while the impact of climate change is controversial, affecting negatively or positively the species distribution, the direction of the effect being species-specific (Téllez-Valdés & Dávila-Aranda, 2003; Martorell *et al*., 2015; Carrillo-Angeles *et al*., 2016).

In recent years species distribution studies have relied on modelling approaches based on powerful statistical techniques and GIS tools and have become popular to predict species distribution, forecast the impact of global climatic change, study species delimitation and niche changes in space or time (Guisan & Zimmermann, 2000; Raxworthy *et al*., 2007; Pearman *et al*., 2008). Niche modelling techniques, of which MaxEnt is the most popular presence-only method, use scenopoietic variables combined to occurrence data to predict the potential geographic distribution and estimate the fundamental niche in the ecological space of the focal species (Soberon & Peterson, 2005; Elith *et al*., 2010). Cacti characterise the landscape of Mexican arid lands and are a significative component of Mexican flora with 68 genera and 689 species, many of which are an important resource, the fruits consumed fresh or dried, the stems being a common food for humans or used as fodder for animals (Casas & Barbera, 2002; Guzmán *et al*.,2007). Although Mexican Cactaceae have a relevant economic and ornamental value, and several studies have analysed their geographical distribution patterns and endemicity (Hernández & Bárcenas, 1995; Hernandez & Barcenas, 1996; Gómez-Hinostrosa & Hernández, 2000; Hernández *et al*., 2001; Hernández *et al*., 2007; Godínez-Alvarez & Ortega-Baes, 2007; Hernández *et al*., 2010), the knowledge of the environmental factors constraining their distribution and characterising their fundamental niche is still quite incomplete, being available only few studies. Cactus richness and endemism in each Mexican state are significantly related to the degree of aridity (Godínez-Alvarez & Ortega-Baes, 2007), a long-term climatic phenomenon, which depends on precipitation, potential evapotranspiration rate, temperature and precipitation seasonality (Maliva & Missimer, 2012), which are the main climatic factors limiting cactus distribution in the Chihuahuan Desert (Hernández & Bárcenas, 1995). Niche modelling has shown that temperature has the greater influence on the distribution of four cactus species in Chihuahua (Cortes *et al*., 2014), while temperature and precipitation are the main environmental variables constraining distribution of *Astrophytum* species (Carrillo-Angeles *et al*., 2016). Temperature influence on cactus distribution can be explained by the low degree of tolerance to freezing temperatures and to the detrimental effect of high temperatures on photosynthetic performance (Flores F & Yeaton, 2003; Aragón-Gastélum *et al*., 2014). This study was aimed at defining the ecological niche of *Thelocactus* species and at quantifying the similarities between them using environmental niche models (ENM) and ordination techniques. Climatic and environmental variables constraining the range of each species were identified and used to generate potential distribution maps. The constructed ENMs were compared to assess the similarities of the ecological niche of the *Thelocactus* species and to know if they share the same ecological niche space.

## Methods

### Studied species and locality data

The nomenclature used is that of Mosco and Zanovello (2000), and nine out of eleven species were included in this study (Table 1). *Thelocactus macdowellii* and *T. lausseri* were excluded because very few populations were available for the analysis, indeed *T. lausseri* is known only from the type locality, while the locality data for *T. macdowellii* were too scanty. Occurrence data were collected from different sources. The majority of them were retrieved from the large dataset build up for the project of mapping Mexican Cactaceae (Hernández & Gómez-Hinostrosa, 2011b), or from the database at the Jardín Botánico Regional de Cadereyta. Additional locality data were obtained from private databases and from personal observations. When only collection site information (state and town name) was available, or when the localities were plotted on 1:250000 topographic maps (INEGI, http://www.inegi.org.mx), their coordinates were inferred using Google Earth. Whenever more records mapped in the same grid cell only one record was retained, as well duplicate records were removed from the dataset.

**Table 1.** Permutation importance of environmental variables to the final Maxent model. The permutation importance represents the drop in training AUC, after the values of the focal variable on training presence and background data are randomly permuted and the model is reevaluated on the permuted data. Permutation values are normalized to percentages.

| Variable | *bicolor* | *buekii* | *conothelos* | *hastifer* | *hexaedrophorus* | *leucacanthus* | *multicephalus* | *rinconensis* | *tulensis* |
| --- | --- | --- | --- | --- | --- | --- | --- | --- | --- |
| Mean diurnal range | 2.4 | 1.8 | 2.5 | 0 | 0.1 | 0.5 | 1.7 | 0.4 | 5.6 |
| Isothermality | 39 | 0.8 | 1.6 | 77.8 | 64.2 | 66 | 33 | 9.8 | 54.4 |
| Max temperature of warmest month | 1.9 | 18.8 | 1.7 | 0 | 1 | 0.4 | 0 | 0.2 | 1.7 |
| Min temperature of coldest month | 12.9 | 7.6 | 0.9 | 6.1 | 5.5 | 5.9 | 24 | 5.4 | 3.6 |
| Temperature annual range | 1.5 | 0 | 3.2 | 0.3 | 3.4 | 0 | 1.2 | 15.9 | 0.4 |
| Precipitation seasonality | 5.3 | 0 | 8.4 | 0.3 | 2 | 17.9 | 8.6 | 0.8 | 1 |
| Precipitation of wettest quarter | 33.1 | 36.6 | 18.8 | 6.1 | 18.7 | 3.8 | 17.2 | 58.4 | 22.6 |
| Precipitation of driest quarter | 1 | 31.4 | 61.2 | 1.1 | 3 | 4 | 6 | 5.6 | 9.6 |
| Aspect | 0.4 | 0.3 | 0 | 0.2 | 0.1 | 0.1 | 1 | 0.6 | 0.2 |
| Terrain ruggedness index | 2.5 | 2.7 | 1.7 | 3.1 | 2 | 1.4 | 7.4 | 3 | 0.9 |

### Predictor variables

Predictor variables were selected from climatic and environmental variables. Climatic variables (precipitation of driest quarter, precipitation of wettest quarter, precipitation seasonality, isothermality, temperature annual range, maximum temperature of warmest month, minimum temperature of coldest month, mean diurnal range) were chosen from the whole set of Worldclim variables (Hijmans *et al*., 2005) after a correlation analysis was conducted on the whole study area, and only variables with pairwise Pearson’s correlation coefficient ≤ 0.7 were retained. The STRM (Shuttle Radar Topography Mission) elevation database aggregated to 30-arc second was the source of digital elevation model (DEM) data (Hijmans *et al*., 2005). Environmental variables, aspect and terrain ruggedness index, were derived from the DEM model using QGIS 2.8.2 software (https://www.qgis.org). Aspect shows the compass bearing of the physical slopes, while the terrain ruggedness index is a quantitative measurement of terrain heterogeneity and was calculated from the DEM according to Riley et al. (Riley *et al*., 1999).

### Species distribution modelling

The study area was confined between 30° N, 107° W and 19° N, 97° W, which represents roughly the distribution limits of the genus. To predict the geographic distribution of *Thelocactus* taxa, models for each species were constructed using the program Maxent (Phillips & Dudík,2008), a machine learning method, which performs well when dealing with presence-only data (Elith *et al*., 2006), even with small datasets (Wisz *et al*., 2008). Maxent was run using its default settings with the number of replicates set to 20. Model performance was evaluated testing the area under curve (AUC) value of the species distribution model (SDM) against a null distribution of 100 repetitions generated by randomly selecting the sample localities from the same geographical space of the studied species (Raes & ter Steege, 2007).

### Niche evaluation

Niche breadth was estimated using ENMTools 1.4.4 (Warren *et al*., 2010) applying the inverse concentration metrics from Levine to the predictions generated by Maxent for each species (Nakazato *et al*., 2010). Niche overlap in ecological space was quantified using an ordination technique that applies kernel smoothers to species densities which ensure that the measured overlap is independent of space resolution (Broennimann *et al*., 2011). The metric *D* was used to calculate niche overlap that can vary from 0, indicating no overlap, to 1, indicating complete overlap (Warren *et al*., 2008). Niche similarity and equivalency tests were conducted in ecological space following the methodology described in Warren (2008). The niche equivalency test was performed to assess whether the ecological niches of *Thelocactus* species are identical. For each species pair, the observed *D* was compared to a null distribution generated by 100 pseudoreplicate datasets. The hypothesis of niche equivalency was rejected when observed values of *D* were significantly (*p* < 0.05) lower than the simulated values. To address whether two environmental niches are more similar than expected by chance, a niche similarity test, based on 100 repetitions, was used. The null hypothesis was rejected if *D* values fell outside the 95% confidence limit of the null distribution.

## Results

### Environmental constraints for Thelocactus species

The distribution of *Thelocactus* species was mostly limited by a combination of two environmental factors, isothermality and precipitation of wettest quarter, but some species were constrained also by other environmental factors (Fig. 1a,b; Table 1). *T. bicolor* is the species with the northernmost distribution, tolerating temperatures below the freezing point, which is reflected by the importance of minimum temperature of the coldest month as a variable along with isothermality and precipitation of wettest quarter. The distribution of *Thelocactus buekii* was constrained by the precipitation of wettest and driest quarter and to a lesser extent by the temperature of warmest month. The precipitation of wettest quarter and of the driest quarter were also the environmental factors of greater importance for the distribution of *Thelocactus conothelos*. Isothermality had the greatest importance, accounting for 77.8% of the total, for *T. hastifer*, as well as for *Thelocactus hexaedrophorus*, whose distribution was constrained, although to a lesser extent, by the precipitation of wettest quarter too. The environmental factors constraining the distribution of *Thelocactus leucacanthus* were isothermality, principally, and secondarily precipitation seasonality. The range of *Thelocactus multicephalus* was limited by three factors, isothermality being the main predictor followed by the minimum temperature of the coldest month and precipitation of wettest quarter. The precipitation of wettest quarter was the main factor constraining the distribution of *Thelocactus rinconensis*, which was limited to a lesser extent by the temperature annual range. Isothermality and precipitation of wettest quarter were the main predictors restricting the range of *Thelocactus tulensis*.

**Figure 1a.**
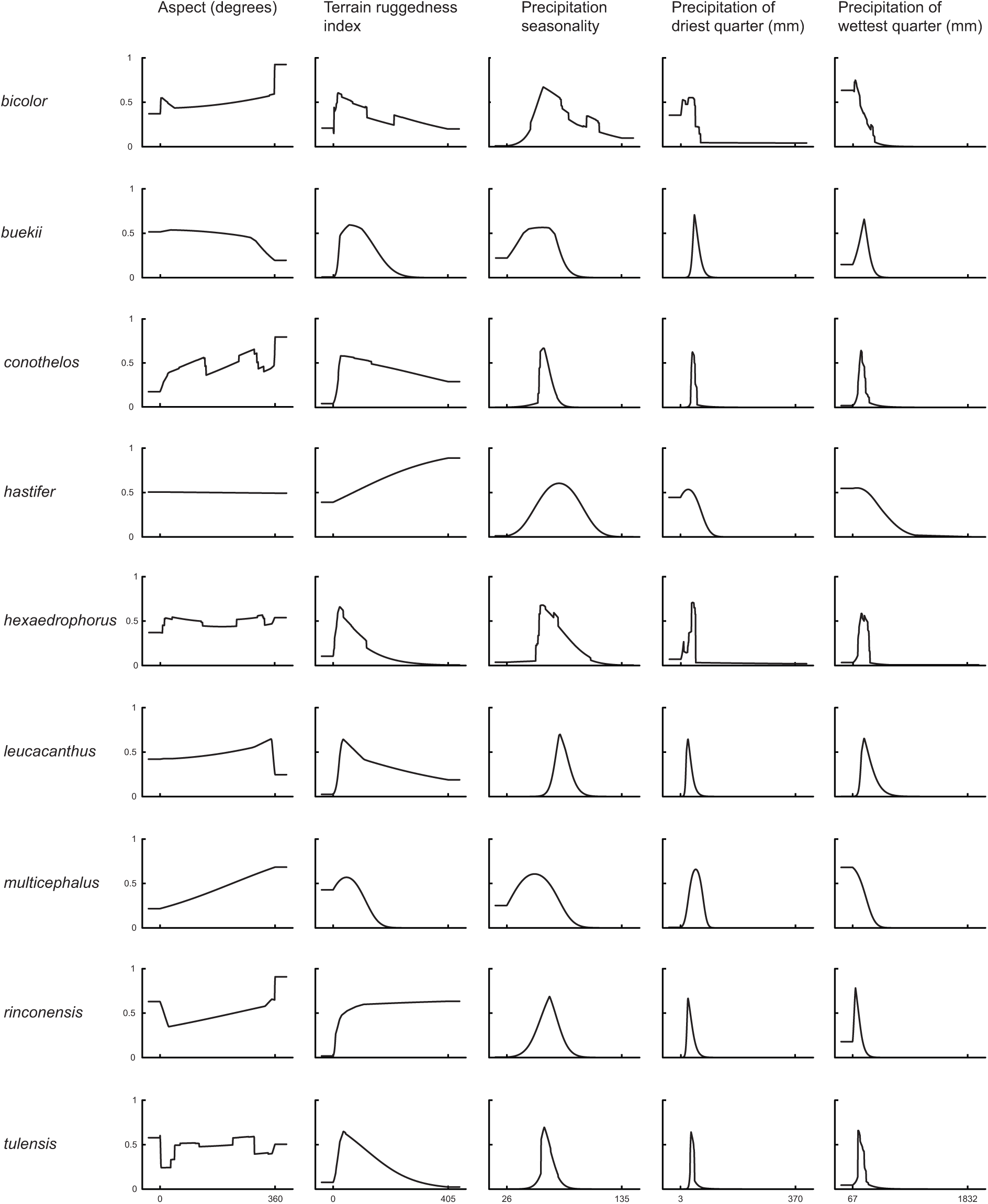
Response curves of Maxent models for *Thelocactus* species. Models were generated using only one variable at a time. The curves show the mean response of the cross-validated models with 20 replicate runs. The value shown on the y-axis is predicted probability of presence.

**Figure 1b.**
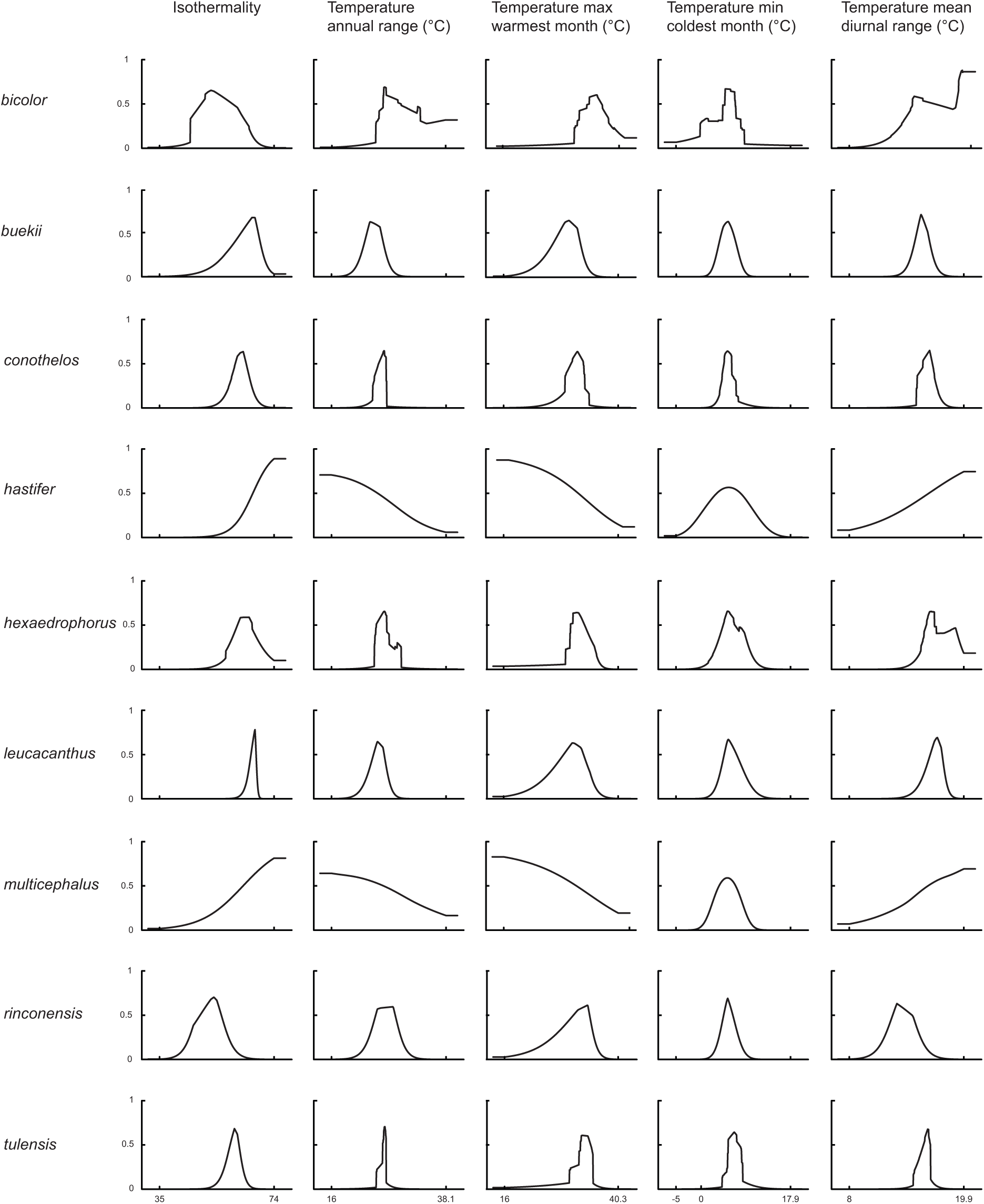
Response curves of Maxent models for *Thelocactus* species. Models were generated using only one variable at a time. The curves show the mean response of the cross-validated models with 20 replicate runs. The value shown on the y-axis is predicted probability of presence.

### Species distribution models

Species distribution models were computed for every taxon. All SDMs, even those generated from few presence records as was the case for *T. hastifer* and *T. multicephalus*, performed significantly better than expected by chance (*p* < 0.001) (Table 2).

**Table 2.** Evaluation of species distribution models by their AUC values, null model results and niche breadth values.

| species | occurrences | AUC | null model AUC | niche breadth |
| --- | --- | --- | --- | --- |
| *bicolor* | 213 | 0.908 | 0.719 | 0.282 |
| *buekii* | 50 | 0.992 | 0.797 | 0.030 |
| *conotheos* | 119 | 0.988 | 0.786 | 0.025 |
| *hastifer* | 11 | 0.969 | 0.847 | 0.178 |
| *hexaedrophorus* | 199 | 0.966 | 0.732 | 0.098 |
| *leucacanthus* | 73 | 0.982 | 0.751 | 0.050 |
| *multicephalus* | 13 | 0.989 | 0.823 | 0.100 |
| *rinconensis* | 47 | 0.978 | 0.792 | 0.062 |
| *tulensis* | 117 | 0.991 | 0.791 | 0.020 |

Most *Thelocactus* species are distributed and have the highest environmental suitability west of the Sierra Madre Oriental (see Appendix S1 in Supporting Information). The predicted suitable areas match well the known distribution of almost all *Thelocactus* species, which confirms that the ecological variables of the models are important for the determination of species’ range. The exception was *T. hastifer*, which has a very small geographic distribution, but much broader suitable areas.

### Ecological niche characteristics

Principal component analysis of environmental niches showed that the first axis explained 34.5% of the total variation, and was mainly loaded by isothermality, the maximum temperature of warmest month, and precipitation seasonality. The second axis explained 27.47% of the variation and was loaded by mean diurnal temperature range and the minimum temperature of coldest month (Fig. 2).

**Figure 2.**
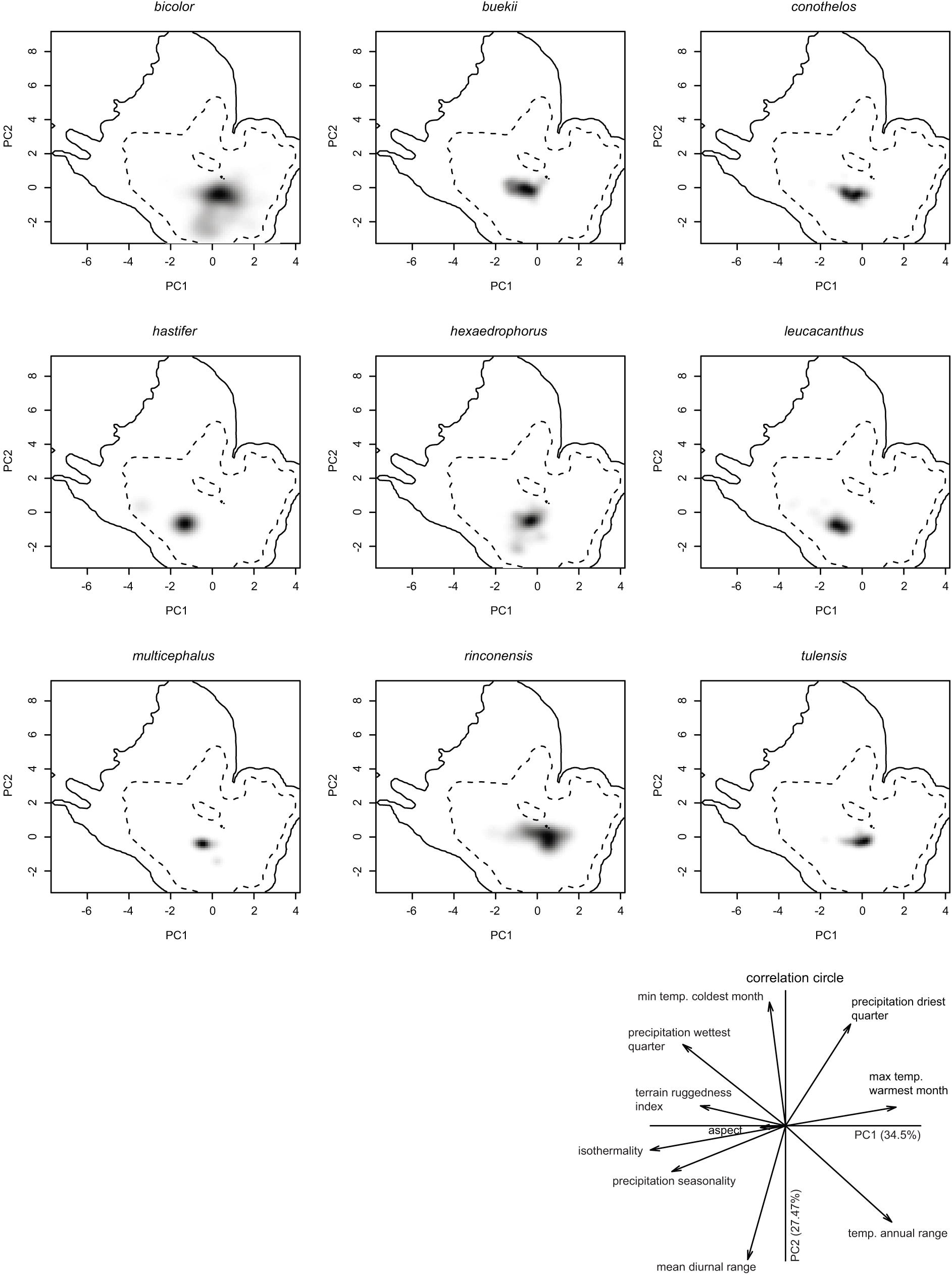
Ecological niche of *Thelocactus* species in environmental space. Niche was displayed in the two main axes of principal component analysis. Grey-to-black shading represents grid cell density of species’ occurrences (black being the highest density). The first dashed line represents the 50% of the available environment and the solid line represents the 100%. The last panel presents the contribution of variables for loading the main PCA-env axes and the percentage of inertia explained by axes one and two.

### Niche breadth and overlap

Niche breadth values varied quite a lot, reflecting the diverse environmental requirements of the studied species (Table 2). *T. bicolor* presented the broadest niche breadth, ten times greater than the niche breadth estimated for *T. buekii, T. conothelos, T. tulensis*, which agrees with its extensive geographic distribution. *T. hexaedrophorus*, which has a broad range, also exhibited a wide niche breadth, although smaller than *T. bicolor*. Another species with a wide niche breadth is *T. multicephalus*, which is proportional to the suitability range and contrasting with the limited known geographic distribution. A case apart is *T. hastifer*, a species known from a very restricted area of about 75-100 km^2^, but with a broad niche breadth, which agrees with the large predicted area of suitability. Niche overlap values are quite variable, ranging from very low values as for *T. leucacanthus* and *T. hastifer* when compared to the other species, indicating that their environmental niches are very different, to higher scores resulting in a partial niche overlap (Table 3).

**Table 3.** Niche overlap values (D) and number of localities where two species coexist. n represents the number of occurrences. In the upper part are displayed the number of localities where two species are simpatric. In the lower part are displayed the niche overlap values (D).

|  | n | *bicolor* | *buekii* | *conothelos* | *hastifer* | *hexaedrophorus* | *leucacanthus* | *multicephalus* | *rinconensis* | *tulensis* |
| --- | --- | --- | --- | --- | --- | --- | --- | --- | --- | --- |
| *bicolor* | 213 | X | 1 | 5 | 0 | 4 | 0 | 1 | 0 | 9 |
| *buekii* | 50 | 0.223 | X | 3 | 0 | 0 | 0 | 2 | 0 | 0 |
| *conothelos* | 119 | 0.203 | 0.539 | X | 0 | 25 | 0 | 2 | 0 | 20 |
| *hastifer* | 11 | 0.056 | 0.158 | 0.193 | X | 0 | 5 | 0 | 0 | 0 |
| *hexaedrophorus* | 199 | 0.344 | 0.506 | 0.644 | 0.192 | X | 0 | 0 | 0 | 15 |
| *leucacanthus* | 73 | 0.068 | 0.177 | 0.254 | 0.626 | 0.257 | X | 0 | 0 | 0 |
| *multicephalus* | 13 | 0.168 | 0.335 | 0.654 | 0.115 | 0.523 | 0.158 | X | 0 | 0 |
| *rinconensis* | 47 | 0.412 | 0.367 | 0.192 | 0.048 | 0.329 | 0.050 | 0.090 | X | 0 |
| *tulensis* | 117 | 0.231 | 0.479 | 0.546 | 0.108 | 0.583 | 0.116 | 0.447 | 0.301 | X |

### Niche similarity and equivalency tests

The null hypothesis of the niche equivalency test was rejected for all paired comparisons between all *Thelocactus* species except between the pair *T. leucacanthus-T. hastifer*. Instead, the results of the niche similarity test were varied. The null hypothesis of niche similarity was rejected for the comparisons between *T. leucacanthus, T. hastifer* and all the other species, as well as for the pair *T. multicephalus-T. rinconensis*. Niche similarity was higher than expected by chance for all other *Thelocactus* species, but for *T. bicolor* compared to *T. buekii, T. hexaedrophorus* and *T. multicephalus*, and *T. rinconensis* compared to *T. buekii, T. conothelos, T. hexaedrophorus and T. tulensis* niche spaces were more similar than expected only in one direction (Table 4). The ecological niche of *T. bicolor* was more similar to those of *T. buekii, T. hexaedrophorus* and *T. multicephalus*, but not vice versa. On the contrary, the ecological niche of *T. rinconensis* was not more similar than expected by chance to those of *T. buekii, T. conothelos, T. hexaedrophorus* and *T. tulensis*, while the null hypothesis held vice versa.

**Table 4.** Comparisons of niche similarity tests. Rows identify the first species of the pairing, columns the second.

|  | *bicolor* | *buekii* | *conothelos* | *hastifer* | *hexaedrophorus* | *leucacanthus* | *multicephalus* | *rinconensis* | *tulensis* |
| --- | --- | --- | --- | --- | --- | --- | --- | --- | --- |
| *bicolor* | X | similar | similar | not significant | similar | not significant | similar | similar | similar |
| *buekii* | not significant | X | similar | not significant | similar | not significant | similar | similar | similar |
| *conothelos* | not significant | similar | X | not significant | similar | not significant | similar | similar | similar |
| *hastifer* | not significant | not significant | not significant | X | not significant | not significant | not significant | not significant | not significant |
| *hexaedrophorus* | not significant | similar | similar | not significant | X | not significant | similar | similar | similar |
| *leucacanthus* | not significant | not significant | not significant | similar | not significant | X | not significant | not significant | not significant |
| *multicephalus* | not significant | similar | similar | not significant | conserved | not significant | X | not significant | similar |
| *rinconensis* | similar | not significant | not significant | not significant | not significant | not significant | not significant | X | not significant |
| *tulensis* | similar | similar | similar | not significant | similar | not significant | similar | similar | X |

## Discussion

The present study has allowed to identify the main environmental constraints that determine the distribution of nine *Thelocactus* species, applying state of the art ENMs and ordination techniques. *Thelocactus* species are mostly endemic or geographically restricted to the Sierra Madre Oriental province, but *T. hexaedrophorus* is present also in the Mexican Plateau province, while *T. bicolor* distribution encompasses several physiographic provinces, extending its range from the Sierra Madre Occidental to the west to the Northern Gulf Plains eastward (Del Conde Juarez *et al*., 2009). At a large scale cactus species richness and endemism in Mexico are mainly related to aridity, while temperature and precipitation are less important, explaining a lower proportion of variance in endemic species (Godínez-Alvarez & Ortega-Baes, 2007). The more detailed analysis of this study shows that the environmental constraints determining the distribution of *Thelocactus* species are more varied.

### Habitat characteristics and environmental constraints

The main factor limiting cactus distribution are sub-zero temperature because they cannot tolerate frost without reporting some damage to their tissues (Gibson & Nobel, 1986). *Thelocactus* species are similarly limited, growing in areas where the temperature does not drop below the freezing point, therefore the minimum temperature of the coldest month is a marginal variable for most of them, with the exception of *T. bicolor* and *T. multicephalus*. For what concerns *T. bicolor*, this is the only species that has a geographic distribution extending northward where freezing temperatures are common, tolerating lower temperatures than the rest of *Thelocactus*, which broadens its range to areas unsuitable to most *Thelocactus* species. Temperature is a limiting factor also for *T. buekii*, but for this species it is important the maximum temperature of the warmest month with predicted suitable areas having maximum temperatures mostly under 30 °C. This reflects the finding of many populations in areas with an arid or semiarid temperate climate with the mean of the maximum temperature of the hottest month not exceeding 22 °C (http://www.inegi.org.mx/geo/contenidos/recnat/clima/default.aspx).

The variable constraining almost all species is the amount of precipitation in the wettest quarter. The Chihuahuan Desert receives low quantities of precipitation, which mostly falls during the summer. The average annual precipitation is 235 mm with a range of about 150-400 mm (Schmidt, 1986). Most cacti occupy areas with dry, semi-hot or semi-dry climates with mean annual precipitations of 300-600 mm, and to a lesser extent dry areas with less than 300 mm per year (Hernández & Bárcenas, 1995). *Thelocactus* species are no exception, occupying areas of low rainfall with a precipitation range that corresponds to that of the response curve of highest probability of suitable conditions generated by Maxent for the precipitation of the wettest quarter. For two species, *T. buekii* and *T. conothelos*, also the precipitation of the driest quarter is important to determine their suitability area, both being more likely to be found in areas with higher rainfall in the driest quarter. The second factor shared by seven out of nine species is isothermality, which quantifies how large are the daily temperature fluctuations relative to annual oscillations. A value of 100 indicates that diurnal temperature range is the same of the annual temperature range, while lower values indicate that day to night temperature oscillations are smaller than annual temperature fluctuations. All *Thelocactus* species are distributed in areas where daily temperature fluctuations are lower than annual ones. The two northernmost species, *T. bicolor* and *T. rinconensis*, have the lowest isothermality index, which agrees with their northern distribution where there is a greater difference between summer and winter temperatures. The potential distribution for *Thelocactus* species obtained from ENMs coincides with the known distribution of most species, suggesting that their distribution is mainly influenced by environmental factors. On the contrary, the potential distribution for *T. hastifer* is much wider than its known distribution. This species is found in a very restricted area of about 75-100 km^2^ at 1800-2000 m with a preference for sedimentary substrates, and its expansion is limited to the south by igneous soils and to the north by the high elevations of the Sierra del Doctor (Sánchez *et al*., 2006). Although it had not been reported from areas north of the Sierra del Doctor, suitable areas are predicted by ENMs to the north and south of the actual distribution. It may well be that more investigation in the most suitable areas could lead to the discovery of other populations (Guisan *et al*., 2006). Indeed, another species growing in the same area of *T. hastifer, Echinocactus grusonii*, was considered to be nearly extinct in the wild after a dam construction led to the loss of its habitat, and only recently a population was found much northerly in Zacatecas (Guadalupe Martínez, Sánchez & Gómez-Hinostrosa, 2013).

### Niche features

The suite of environmental conditions or resources that a species can inhabit or use describes its niche breadth (Gaston *et al*., 1997), the larger the niche breadth the wider the environment spectrum a species utilises. *T. bicolor* has the broadest niche breadth as well as the largest distribution range and therefore is tolerant of a wider spectrum of climatic conditions that promoted a large morphological variation among local populations, some of them being recognised at subspecific level. At the northern border of the Chihuahuan Desert, we find *T. bicolor* ssp. *flavidispinus* which is endemic to caballos novaculite outcrops in Brewster Co., Texas. In Coahuila, *T. bicolor* ssp. *bolaensis*, an ecotype with white spines, colonises the limestone slopes of the Sierra Bola. To the west, in Zacatecas and Durango *T. bicolor* ssp. *heterochromus* is found, while at the eastern range of the species distribution we find *T. bicolor* ssp. *schwarzii*, which is the only *Thelocactus* growing outside the Chihuahuan Desert, in Tamaulipas, on rocky outcrops in the Tamaulipan thorn shrub. The large niche breadth, which is related to the amount of environmental variation, makes this species a generalist, adapted to different environmental conditions (Kassen, 2002; Dennis *et al*˙., 2011). The second widest niche breadth is that of *T. hastifer*, which is in agreement with its large potential distribution range, but contrasts with its very restricted geographical range. This discrepancy can be explained by the small number of occurrences, distributed in a very restricted area, although was shown that Maxent performs well also with small sample sizes, or be caused by factors different from climatic predictors and as such missing from the model (Wisz *et al*., 2008; Nakazato *et al*., 2010). The other species display smaller niche breadths and, with the exception of *T. hexaedrophorus*, also small geographical ranges. These results indicate they are less tolerant toward a wide spectrum of climatic variation, preferring more homogeneous environments, which is typical for specialist organisms (Kassen, 2002; Dennis *et al*., 2011). Niche overlap values between *Thelocactus* species are mostly low, reflecting the difference in the environmental suits each species is adapted to. With the exception of the pair *hastifer-leucacanthus*, the niche equivalency test was rejected for all other species, showing that environmental spaces of *Thelocactus* species are significantly different from each other (Warren *et al*., 2010). The results of the niche similarity tests were quite varied, for some species pairs the niche similarities were higher than expected by chance, for others the null hypothesis was rejected, while in other species pairs niches were more similar than expected by chance, but only in one direction. The niches of the pair *T. hastifer* and *T. leucacanthus* were more similar than expected by chance, which was expected as their niches are equivalent, while when compared to the other species the similarity was not significative. Considering that these two species have also an overlapping geographical distribution, the results obtained support the hypothesis of niche conservatism. Niche similarity higher than expected by chance was found also in most of the other pair-wise comparisons, suggesting that habitat conservatism is common among *Thelocactus* species. *T. bicolor* has the widest niche breadth, therefore being capable to exploit a larger set of environmental conditions, which is in agreement with its large geographic distribution that overlaps with the range of most species, with the exclusion of the two southernmost, which is probably the explanation for the background test being significantly more similar when compared to the other species. The reverse was not always true. For *T. buekii, T. conothelos, T. hexaedrophorus* and *T. multicephalus* the results were not significantly similar, suggesting that these species are not suited to the habitat conditions in which *T. bicolor* can grow. This result is probably a consequence of the high heterogeneity of *T. bicolor* habitat (Nakazato *et al*., 2010). For what concerns *T. rinconensis* the similarity test was rejected when paired to *T. buekii, T. conothelos, T. hexaedrophorus* and *T. tulensis*, but the reverse comparisons showed that the background test was accepted. These results suggest that *T. rinconensis* has rather different environmental requirements of the other four species and that these have a more heterogeneous habitat and therefore their niches overlap that of *T. rinconensis*. Five species, *T. buekii, T. conothelos, T. hexaedrophorus, T. multicephalus* and *T. tulensis*, showed a similarity greater than expected by chance. All of them are geographically distributed in part or only in the Galeana, Mier y Noriega and Huizache subregions of the CDR (Hernandez & Barcenas, 1996), areas rich in species number and endemicity of Cacteae, whose diversification is related to increased aridity in response to the uplift of the Sierra Madre Oriental and the development of the Trans-Mexican Volcanic Belt (Vázquez-Sánchez *et al*., 2013) in the late Miocene (Arakaki *et al*., 2011; Hernández-Hernández *et al*., 2014). Pleistocene glacial maximum (Wisconsin, 11000 years ago) brought a cooler and wetter climate affecting the areas occupied by desert communities. Climate fluctuations driven by advances and retreats of the Laurentide Continental Glacier promoted contractions, retractions and displacements of the geographic range of the species involved (Cartron *et al*., 2005). It was suggested that the richest areas in cactus taxa in the CDR acted as glacial refugia, leading to isolation and species diversification as well as shaping actual cactus distribution (Hernández & Bárcenas, 1995; Gómez-Hinostrosa & Hernández, 2000). Although the observed geographic ranges of *Thelocactus* species rarely overlap, and then mostly partially, and that species distribution is mainly allopatric, significant ecological niche conservatism is found for most species pairs, indicating that many *Thelocactus* species have conserved their ecological niche traits over time (Wiens *et al*., 2010), while the significant results of the background test in only one direction, and not significant in the counter-direction, involving some species pairs, probably depend on the differences in the environmental background for the species pairs (Nakazato *et al*., 2010; Theodoridis *et al*., 2013).

### Climatic change and conservation issues

Cacti are the fifth most threatened major taxonomic group with 31% of species threatened. Land conversion to agriculture affects large parts of cactus species in Northern Mexico, while the unscrupulous collection of plants and seeds is the main risk factor for threatened cacti (Goettsch *et al*., 2015). Nevertheless, future climate change may play an important role in redesigning distribution ranges of current populations, in the worst case leading to extinction (Téllez-Valdés & Dávila-Aranda, 2003; Martorell *et al*., 2015). Projected climates for the following years show an increase of the mean annual temperature by 1.5 °C in the decade around 2030 and a decrease in precipitation, with an expansion of the arid zones of north-central Mexico toward both coasts and south-east (Sáenz-Romero *et al*., 2010). Although in general cacti should benefit from an increase in CO_2_ concentration and temperature rise, extending their poleward and elevation ranges, the impact of climate change should be determined at specific level (Nobel, 1996) (Munson *et al*., 2012). Indeed, niche projections for future climate show that species would respond in specific ways, the predicted distribution areas varying from remaining stable to undergoing a severe contraction (Aragón-Gastélum *et al*., 2014; Cortes *et al*., 2014; Carrillo-Angeles *et al*., 2016). However, the potential distribution areas may not match potentially colonizable areas, the process being limited by several factors as seed dispersal efficiency, spatial barriers and unconnected distribution areas. The seed dispersal mechanism for *Thelocactus* species has a low efficiency because seeds are released from dry fruits by a basal pore and then fall on the ground. The presence in many *Thelocactus* species of areolar glands secreting a sweet nectar may serve to attract ants favouring seed collection and subsequent transportation. Myrmecochory is probably the most frequent mechanism of seed dispersal in Cactaceae, but it does not allow the reach of great dispersal distances (Bregman, 1988; Vittoz & Engler, 2007). Given these limitations coupled with a slow growth rate and a limited plant recruitment caused by seedling sensitivity to high temperatures (Aragón-Gastélum *et al*., 2016), makes *Thelocactus* species highly vulnerable to climate warming. Chihuahuan Desert hosts several protected areas (Bezaury-Creel *et al*., 2007), both at federal and state level, and most *Thelocactus* species can be found in some of them (see Appendix S1 in Supporting Information), although the percentage of localities occurring in protected areas is generally low (Hernández & Gómez-Hinostrosa, 2011a). The situation is worse for microendemic taxa that occur in very small areas, e.g. some *T. bicolor* and *T. conothelos* subspecies and *T. hastifer*, which do not occur in any protected area and for which the creation of small reserve areas was already proposed in view of its efficacy and as a complement to largest protected areas (Anderson *et al*., 1994; Hernández & Gómez-Hinostrosa, 2011a).

## Conclusions

The present study has assessed the characteristics of the fundamental niche of most *Thelocactus* species, their niche breadth and their potential distribution range. The spatial predictions could serve to improve field design sampling to discover new populations, above all for the most threatened species, an effort at the base for the delimitation of potential new reserve areas. Differences in habitat requirements are well documented by the significative differences in the niche ecological space as shown by the equivalency test, while the high percentage of niches that are more similar than expected by chance suggest a high degree of niche conservatism among *Thelocactus* species, which is supported by the close geographic distribution areas of many species, above all in the Galeana, Mier y Noriega and Huizache subregions of the CDR, or the overlapping distribution in the southernmost part of the CDR (see Appendix S1 in Supporting Information). Several threats to Cactaceae have been recognised, of which global warming can impact many species acting directly on their physiology and ability to adapt to new environmental conditions or indirectly changing the biotic interactions, as climate change could affect also the presence of pollinators and the animals required for seed dispersal or have an effect on the vegetation community and the nurse plants belonging to it (Ibisch & Mutke 2015). The necessity of increasing protected areas in the CDR has been already underlined and the results presented in this study could be relevant for improving preservation actions and guiding reintroduction programs for a better conservation of *Thelocactus* species, taking into account the ecological requirements of focal species.

## Acknowledgements

I am thankful to Héctor Manuel Hernández, Carlos Gómez Hinostrosa, Johann Jauernig, and Emiliano Sánchez Martínez, who provided a great part of the data for *Thelocactus* localities.

